# Ancient genomes from the siege and destruction of Middle Bronze Age Roca Vecchia (Apulia, Italy) shed light on Aegean contacts and conflicts

**DOI:** 10.64898/2025.12.15.694319

**Authors:** Serena Aneli, Valeria Nicolini, Giorgia Vincenti, Francesco Montinaro, Stefania Sasso, Tina Saupe, Helja Kabral, Anu Solnik, Kristiina Tambets, Riccardo Guglielmino, Pier Francesco Fabbri, Luca Pagani

**Affiliations:** Department of Public Health Science and Pediatrics, University of Turin; Department of Biology and Biotechnology “L. Spallanzani”, University of Pavia; Physical Anthropology Laboratory, University of Salento; Department of Biology, University of Bari; Institute of Genomics, University of Tartu; Human Evolution, Department of Organismal Biology, Evolutionary Biology Centre, Uppsala University, Uppsala, Sweden; Museo Fiorentino di Preistoria; Department of Biology, University of Padova

## Abstract

**Background:** Roca Vecchia, an iconic Bronze Age stronghold in Apulia, Southern Italy, was completely destroyed during a siege between the end of the 15th century BCE and the beginning of the 14th century BCE. During the siege, seven of the local people hid within the stronghold walls. Two others, who could have been as well attackers as defenders, were found under the ruins of the main gate. The material culture found at Roca Vecchia and associated with the period of the siege includes Minoan-type pottery produced from local clay, imported Aegean pottery and an Aegean-type dagger, pointing to an established relationship between the site and the Minoan civilization. Therefore, the site offers an unprecedented opportunity to characterise the genetic components of the population inhabiting an indigenous settlement with increasing contacts with the Aegean world, and to shed light on the demic or cultural modes of the Minoan presence in the central Mediterranean.

**Results:** With our work, we sampled six out of nine available unburied Middle Bronze Age individuals, contemporary with the siege and destruction of the site, and obtained genome-wide information for two individuals. When compared with available Minoan, Apulian and broadly Mediterranean genomes, the individuals showed a characteristic Bronze/Iron Age Italian peninsula genetic signature, with limited contribution from Minoans.

**Conclusions:** We conclude that the local population of Roca Vecchia, at the moment of the siege, was predominantly autochthonous, with a minoritarian Minoan component. A Minoan genetic signal is indeed likely present in one out of two analysed individuals who were certainly part of the dwellers of Roca Vecchia. This confirms previous hypotheses supposing that a nucleus of “foreigners” coming from the Minoan world was living in the site and mixed with locals. Archaeological data suggest that the Roca Vecchia Aegean population component probably increased in the following centuries.

## Background

Roca Vecchia (Melendugno, Lecce), located along the Adriatic coast, is one of the most important Middle Bronze Age long-term fortified sites in Italy (Figure 1A). During the Middle Bronze Age (MBA), this site established strong trade connections with the Aegean world [1]. In particular, the finding of local pottery resembling Aegean styles led to the suggestion that ceramists from Crete or mainland Greece may have lived in Roca Vecchia during the MBA, as a well-defined community [2–4].

**Figure 1.**
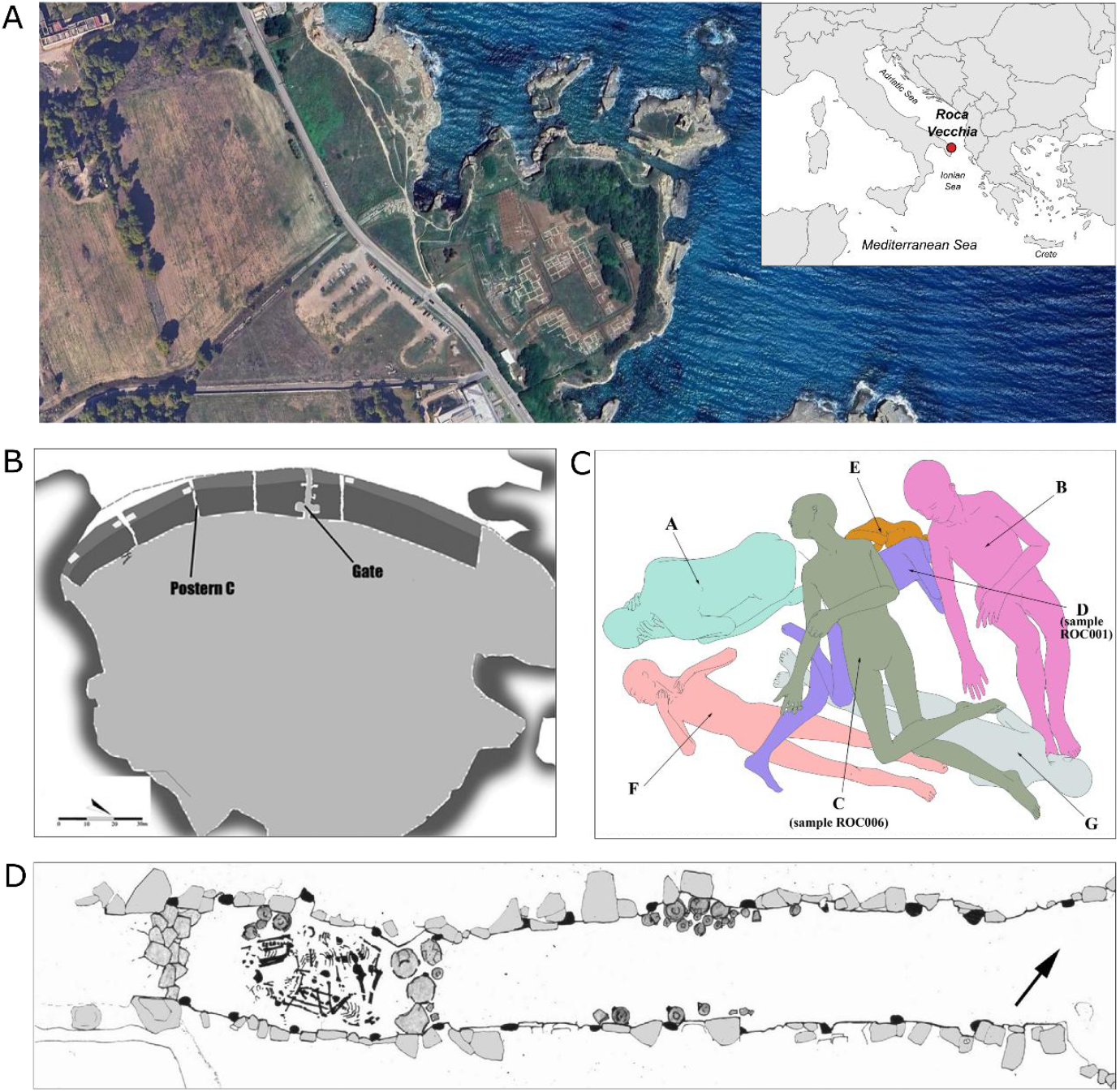
The Middle Bronze Age site of Roca Vecchia, Southeastern Italy. A and inset) Map of the archaeological site. B) The Middle Bronze Age fortification. C) Representation of the individuals found in Postern C of the fortifications. D) Schematic representation of Postern C and the location of seven skeletons investigated in this study.

Roca Vecchia was destroyed by a siege-induced fire, which caused the collapse of the fortifications [1,5] during Middle Bronze Age 3 (MBA3), specifically between the second half of the 15th century BCE and the first decades of the 14th century BCE (3545-3331 calBP, 95% CI). This chronology is based on statistical analysis of radiocarbon dates from short-lived plant remains recovered from different parts of the collapsed structures, together with stratigraphic data [6].

Excavations of the fortifications revealed the remains of at least ten individuals (see Supplementary Note in the Supplementary Materials). Seven of them, labelled US 2616 A to G, were in postern C of the fortifications (Figures 1B-D, Supplementary Figures 1 and 2). They included two adults (US 2616 A and US 2616 B, one male and one female, based on osteological evidence) and five subadults ranging from 4 to 16 years old (an adolescent labelled US 2616 C and four children labelled US 2616 D-G) [7]. Another individual (US 14203 RA7, Supplementary Figures 3 and 4), an adult male not included in this study, was found near the interior entrance of the main gate during the excavations conducted by the Superintendence of Lecce [8]. A young male skeleton (US 813 RA2, Supplementary Figures 5 and 6), who was stabbed to death, was instead found in room B of the main gate [9].

**Figure 2.**
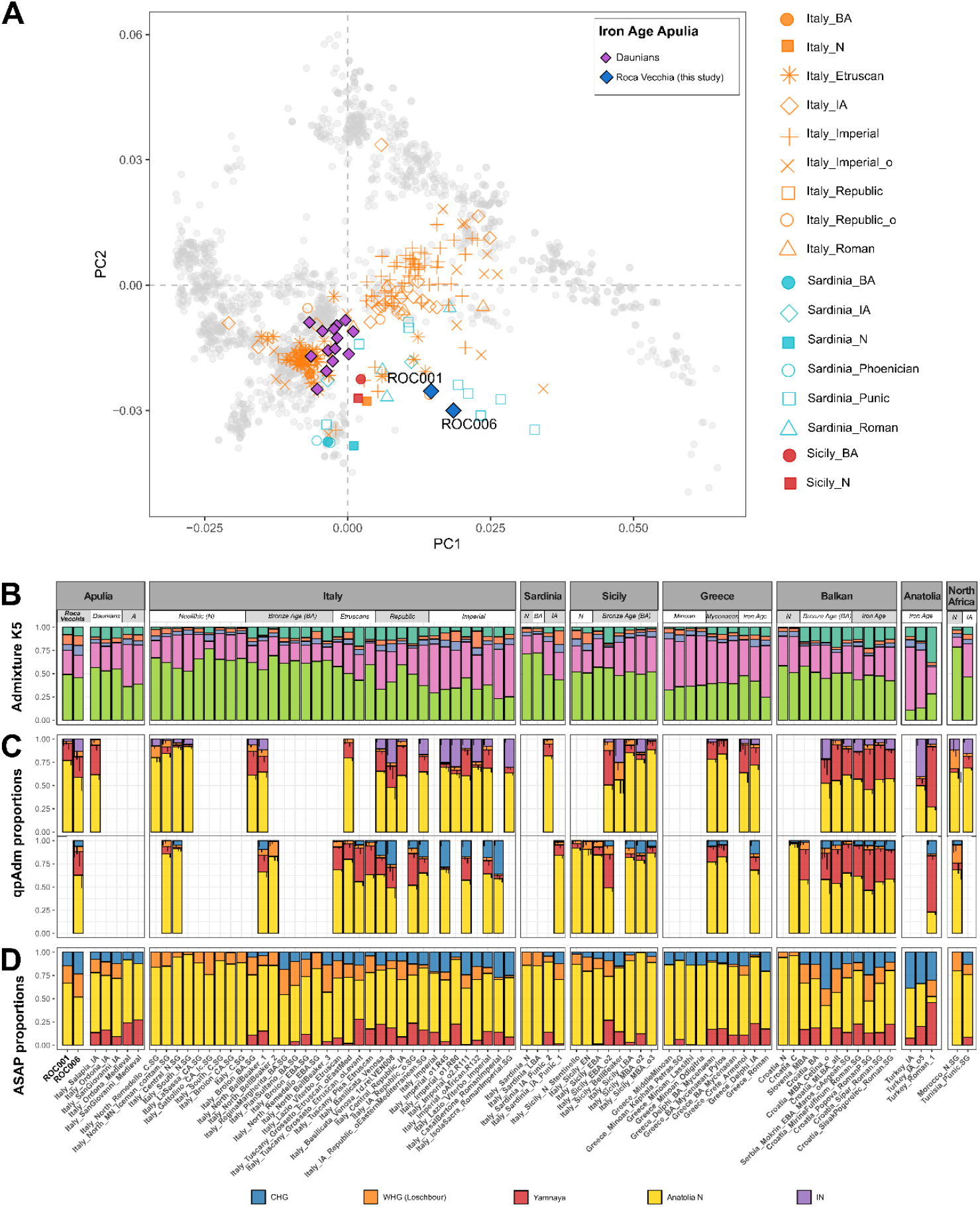
Genetic description of ROC001 and ROC006 samples within the context of available ancient DNA from Europe and the Mediterranean region. **(A)** Principal component analysis (for the Neolithic and Bronze Age samples, only average positions are plotted). More reference samples from Europe and the Mediterranean basin are shown in Supplementary Figure 8; **(B)** projected Admixture (for colour reference, see Supplementary Figure 8); **(C)** qpAdm models with Iranian Neolithic (top) or CHG (bottom) as alternative sources. Just the models with non-negative admixture proportions and a p-value greater than 0.05 are shown (the complete list of models is in Supplementary Tables 6 and 7). **(D)** PANE ancestry composition. N: Neolithic, BA: Bronze Age, IA: Iron Age, A: Antiquity.

Archaeological evidence indicates that the posterns had been transformed into shelters during the siege. Indeed, a great amount of pottery, similar to that usually retrieved in dwelling places, was found within them. Moreover, the presence in the fortifications of subadults and unarmed individuals associated with domestic pottery suggests that civilians sought refuge there because of the state of emergency. Conditions similar to those observed in Roca are mentioned in various historical literary sources. For example, Thucydides states that, in Athens during the Peloponnesian War, many refugees from the surrounding areas sheltered in the fortification towers of the Long Walls and the Walls of Piraeus (Thuc., 2, 17). The reuse of posterns as dwelling areas is probably due to the overpopulation of the sites during war times, when people living in the surroundings found shelter inside the fortification; a similar behaviour has been hypothesized for the Mycenaean citadels and Troy [10,11].

Postern C, the main subject of this study, was filled with a thick layer, labelled US2616, consisting of burned stones, ashes, and charcoals from the collapse of overhanging fortification structures. At its base lay the seven skeletons mentioned above (Figure 1C), in a confined space of about 2×1.5m bordered by the wall obstructing the outer entrance and a row of large vases. The skeletons were partially entangled, the articulations were nearly perfectly preserved, and the individuals at the bottom of the deposit had not been displaced by those on top. Based on these observations, the simultaneity of the deposition of the seven individuals [12] before the destruction of the fortification can be safely argued. In addition to pressure-induced breakings, heat-induced chromatic modifications or destruction of bones are common, but very variable in intensity. On the same individual, often on the same bone, we can observe the full range of fire-induced modifications: from unaffected to completely charred. Although scattered human remains and graves have been found in some Bronze Age sites [13], postern C cannot be interpreted as a grave for several reasons: it is not similar to MBA graves found in the region [14]; the structure, where the skeletons have been found, was part of the fortification and was used as a dwelling structure at the moment of its destruction [15]; the big vases around the skeletons are not usually found in graves where instead small and probably ad hoc made vases are common [16]; bodies had not been composed in the ritual position that is laying on a side with flexed upper limbs [14]. From the archaeological examination of the site (for further details, see Supplementary Note), one can therefore conclude that, in all likelihood, the individuals found at the postern C can be safely interpreted as part of the local population rather than as part of the people who promoted the siege. In addition, Sr isotope analyses on both US 2616 D and US 2616 C are consistent with a local, coastal geographic origin [9].

With the present study, we aim to investigate the genetic origin of six of the individuals found under the debris of the MBA fortifications: five non-belligerent inhabitants from postern C (US 2616 A, C, D, E and G) and one young male from room B of the main gate (US 813 RA2) who was a belligerent, potentially either a defender or an attacker, and died in combat. Because the individuals recovered from postern C were part of the local civilian populations, they provide a unique opportunity to investigate the demographic history of this iconic Southern Italian Bronze Age stronghold. At the same time, the distinctive Aegean-style ceramics found at Roca Vecchia raise the question of whether this cultural influence corresponded to actual population movements [2–4]. Our goal, therefore, is also to explore possible genetic links with the Aegean world, which may elucidate whether an Aegean demic component was present in Roca Vecchia.

## Results

We extracted and sequenced DNA from 6 selected individuals, labelled ROC001 to ROC006. Five corresponded to those with the least evidence of heat damage from Postern C, while the sixth referred to another individual recovered at the site in a chronologically consistent layer (Table 1 and Supplementary Table 1). Radiocarbon dating was attempted as part of a previous study but none of the human samples yielded sufficient information. However, the clear stratigraphic association between the human remains and the collapsed ruins enables us to indirectly date the analysed human remains by exploiting C14 dates available from short-lived plants found within the strata (3545-3331 calBP, 95% CI) [6].

**Table 1.**
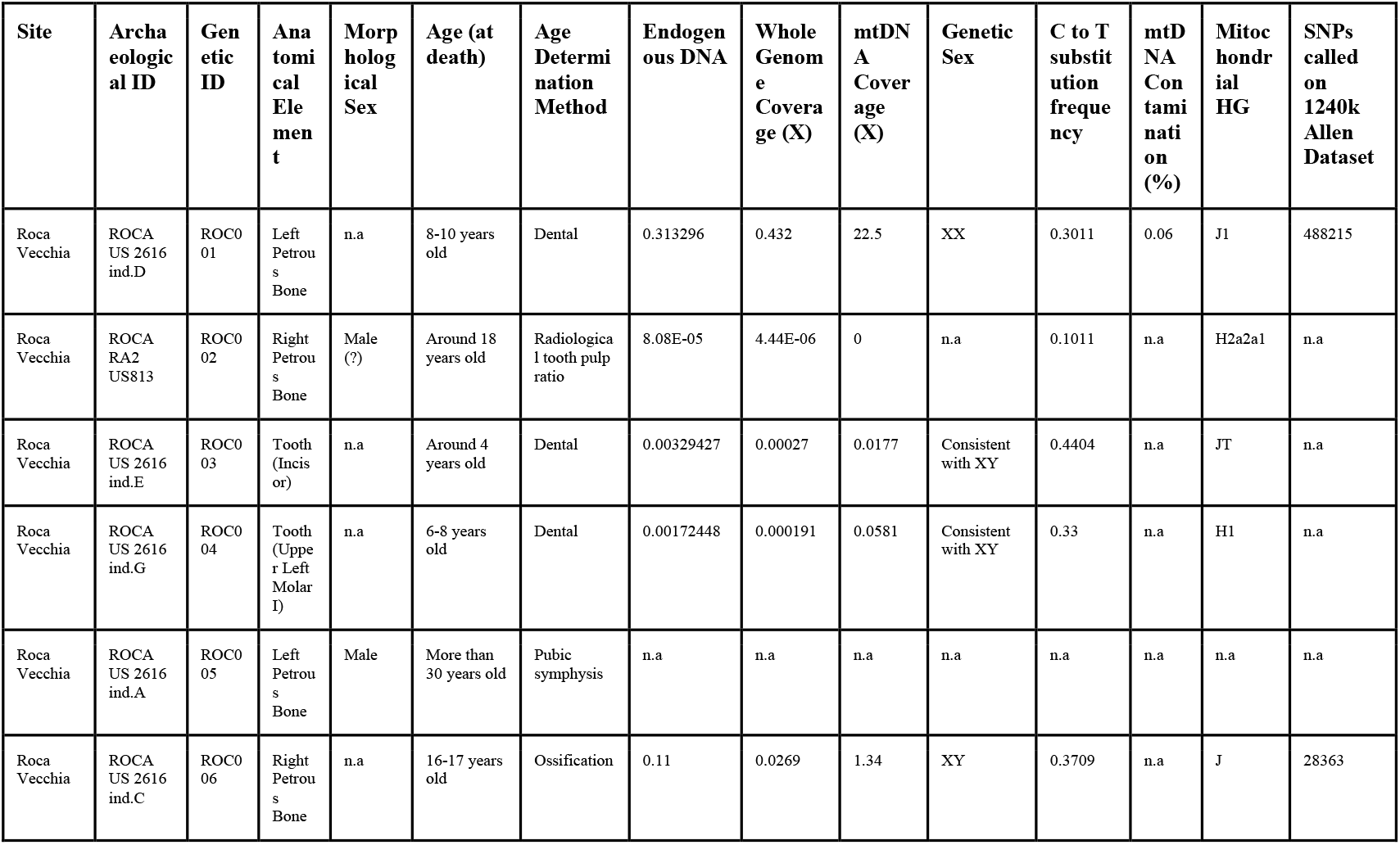
Roca Vecchia individuals for whom DNA sequencing was attempted in this study. mtDNA - mitochondrial DNA. NA - not available. Coverage measurements refer to sequencing depth. Further details can be found in Supplementary Table 1.

Despite multiple extraction attempts, we could retrieve human DNA only from two individuals from Postern C: a child (genetic ID: ROC001 archaeo ID: ROCA US 2616 ind.D) and a non-adult (genetic ID: ROC006 archaeo ID: ROCA US 2616 ind.C). The low success rate could be explained by the intense heat to which the bones were exposed during the fire that followed the siege, although the specimens did not show typical signs of combustion. aDNA libraries were prepared using a non-UDG protocol. We then used MapDamage2.0 [17] to estimate cytosine deamination rates and the resulting C to T substitution at the 5’ termini of DNA fragments, which are typically due to sample age and environmental conditions. Substitution rates were estimated at 30% for ROC001 and 37% for ROC006 (Supplementary Figure 7).

Genetic sex was determined for ROC001 and ROC006, which are XX and XY, respectively. We could confidently call 488,215 SNPs for ROC001, and 28,363 for ROC006, among the ones listed in the standard 1240K SNP set described in the Allen Ancient DNA Resource. We also estimated mitochondrial DNA to be less than 5% in ROC001, while, given the low coverage of ROC006, no contamination estimate was possible. Genetic relatedness between ROC001 and ROC006 was then assessed. Due to low coverage and the limited number of samples in our dataset, kinship analysis was performed both with and without a normalization value, computed on a genetically similar reference population [18]. Without normalization, kinship values did not indicate biological relatedness. Applying the normalization value yielded consistent results, confirming the absence of detectable genetic kinship (Supplementary Table 2).

To provide an overview of the genetic variation of the two Roca Vecchia individuals, we performed a Principal Component Analysis (PCA) in the context of West Eurasian genetic variation (Figure 2A, Supplementary Figure 8, Supplementary Tables 3 and 4). At first glance, the two Roca Vecchia genomes occupy a peculiar place within the PCA space obtained after projecting ancient samples onto the present-day West Eurasian genetic variability. The two samples are indeed outside of the cloud defined by Daunian samples [18], the Iron Age population that inhabited the Apulian area some 500-1000 years after the siege (after the 10th Century BCE), and they are also departed from any Minoan or Mycenaean sample available to date. This preliminary observation may therefore point to a lack of substantial genetic contribution from Aegean populations to the genetic makeup of the site, as well as a not obvious continuity between the Bronze Age and the subsequent Iron Age Apulian groups. To further validate the location of the two Roca Vecchia individuals within the PCA space, particularly ROC006, which has a lower number of called SNPs (Table 1 and Supplementary Table 1), we performed an additional PCA using the full 1240K SNP set (see Methods, Supplementary Figures 9, 10 and Supplementary Table 4). This approach maximizes SNP overlap between ancient and modern samples and confirms the relative positioning of ROC006, which remains broadly consistent despite its lower genomic coverage.

At a finer scale, however, the formal tests of allele frequency correlations between populations of *f*4-statistics [19] in the form of *f*4 (Bronze Age Roca Vecchia; Iron Age Apulia; X; Mbuti) could not differentiate RocaVecchia from subsequent Iron Age Apulian populations, with X being a Bronze Age Mediterranean population or a distal population source such as Anatolia_N (Anatolian Neolithic); CHG (Caucasian Hunter-Gatherers); Iran_N (Iranian Neolithic); Yamnaya (Steppe-related ancestry) or North African (as a putative additional contributor to the Mediterranean genetic makeup) (the full list of f4 analyses and their rationale are in Supplementary Table 5, while the complete list of individuals used for such analyses is in Supplementary Table 3). We therefore resorted to Admixture [20] and qpAdm [19] to model the genetic composition of these ancient individuals, and to delve into the apparent differences that emerged from our preliminary PCA analyses.

Admixture analysis, obtained via projecting the ancient genotypes onto the allele frequency space obtained from present-day individuals, highlighted an overall similar genetic composition of Roca Vecchia and Daunian samples, with a slight affinity of the former to ancient and present-day North African and Levantine individuals (Figure 2B and more in details in Supplementary Figure 11), which as mentioned was not confirmed by *f*4 tests (Supplementary Table 5).

We then modelled Roca Vecchia, Daunian, and other Bronze Age Mediterranean individuals using distal sources known to have contributed genetic components to the area during post-Neolithic movements (Supplementary Tables 6-9) [21]. We report for both Roca Vecchia individuals an admixture profile typical of Bronze Age Italians [22], with their genomes being constituted predominantly by Anatolian_N and WHG (Western Hunter-Gatherers) components (Figure 2C), the latter being a diagnostic constituent of Mainland European populations, as opposed to Aegean groups of the same period. However, we noted that while ROC006 has a similar genetic composition to the subsequent Salapia and Ordona Daunian groups, ROC001 shows an increased Anatolia_N component, matched by reduced WHG and CHG components (Figure 2C and Supplementary Tables 6 and 7). This difference, albeit small, may be compatible with a rather more Aegean contribution in the genome of the ROC001 child. The genome of ROC001 can indeed be modelled as roughly 50% of ROC006 (or Salapia or Ordona) and 50% of Minoan when taking linear combinations of their ancestry composition (available from Supplementary Table 10). This genetic picture was confirmed using PANE [23], a newly developed method that leverages the first 100 principal components to model a target genome as a linear combination of a pre-defined set of source individuals via non-negative least squares (NNLS) optimization (Figure 2D and Supplementary Table 10).

To further dissect the contribution of Steppe-related ancestry, we applied two additional qpAdm frameworks (see Methods and Supplementary Tables 8 and 9). A distal model including Anatolia_N and three hunter-gatherer components (WHG, CHG, and EHG: Eastern Hunter-Gatherers) successfully modelled ROC001, confirming the presence of a ∼15% Steppe signature (1.8% CHG and 13% EHG), albeit with non-negligible standard errors (Supplementary Table 8). A proximal model, instead, used a European Chalcolithic group lacking Steppe ancestry together with Yamnaya as sources (Supplementary Table 9). Under this framework, only ROC006 could be successfully modelled (76% European Chalcolithic, 24% Yamnaya), supporting the presence of the Steppe-related ancestry in Apulia, at the Southern tip of the Italian peninsula, by ∼1500 BCE, contemporaneously with central Italy [22]. The successful fit of ROC006 with the proximal model further suggests that this individual represents a more autochthonous genetic profile, in line with local Bronze Age Italians. By contrast, ROC001 could not be modelled under this scenario, as it harbours a reduced proportion of WHG ancestry (present in the European Chalcolithic source) compared to ROC006, pointing instead to stronger affinities with Aegean populations rather than with local Bronze Age Italians.

When looking at finer details of the emerging scenario, we also report that both qpAdm and PANE inferred for the individual ROC006 a sizeable amount of CHG (Figure 2C and D, blue) and/or Iranian Neolithic components, with a magnitude that is matched only by Imperial Roman or even more recent samples, with the exception of one Bell Beaker sample from Sicily. qpAdm also detected a Steppe-related component in ROC006 (Figure 2C, red), which was not replicated by PANE (Figure 2D), perhaps due to the overall similarity between CHG and Steppe-related components that may have confounded one of the two analyses also in the light of the low sequencing coverage achieved for both samples.

## Discussion

With this study, we provide a characterization of the genetic makeup of local individuals from Roca Vecchia, a Middle Bronze Age fortified site in coastal Southern Apulia, that fell under a siege between the second half of the 15th century BCE and the first decades of the 14th century BCE and was eventually burnt and destroyed. On top of an autochthonous Italian signature, represented by the Anatolian Neolithic and WHG genetic components, the sequenced Bronze Age samples show a clear discontinuity with any Italian Neolithic samples sequenced so far, displaying a prominent genetic signature that can be ascribed to Iran Neolithic or CHG ancestries, putatively coming from the Levant and which make the analysed Bronze Age Apulians genetically similar to the Iron Age populations that gave birth to the Daunian civilization. In particular, we here report the earliest signature of CHG/Iran Neolithic component in continental Italy, dated to at least 3500 years ago. In light of the lesser extent of CHG/Iran_N in many Iron Age Daunian samples in Apulia, we hypothesize a dilution of such a signature, perhaps in favour of more Anatolia_N components coming from the Mediterranean area. Our qpAdm results also show that the Steppe genetic signature had reached as far south as Apulia at the same time when it first appeared in Veneto (Broion shows Steppe 3500 years ago and not earlier [22]) and Remedello. On the other hand, the same signature could not be replicated with the recently introduced PANE method. Besides the overall low sequencing coverage achieved for the Roca Vecchia samples, a reason for this discrepancy may be due to the partial similarity between Steppe and CHG components, which may result in mimicking each other in this kind of genetic inferences. As far as the CHG signature is concerned, however, we are confident about its presence given that it was possible to model it with qpAdm, allowing at the same time for the presence of the Steppe component. On the other hand, no direct North African contribution to the Rocavecchia samples could be formally confirmed by f-statistics, despite the suggestive right-shifted position of these individuals in the PCA and the presence of a minor ancestry component in unsupervised ADMIXTURE analyses. However, given the limited resolution of the data and the sensitivity of such tests to outgroup choice, a subtle African-related contribution cannot be excluded and will require further investigation. An interesting feature emerging from the analysis of our newly sequenced samples, within the context of other data available from the literature, is that during the Bronze Age, the Italian population already showed a sizeable fragmentation, with Italian Bronze Age samples spanning a gradient from Sardinia to Roca Vecchia in the PCA.

## Conclusions

In conclusion, we show that the genetic composition of ROC006 (US2616 C), an adolescent male aged 16-17 years, is too divergent from the ones displayed by available Greek samples from a comparable period, pointing to a local rather than Aegean origin for this individual, who can be safely described as a pre-Daunian, with a slightly higher fraction of CHG/Iran_N ancestry. On the other hand, ROC001 (US2616 D), a female child aged 8-10 years, may be of similar genetic makeup or a mix between such a signature and a more Aegean genetic makeup, given the observed lower WHG and higher Anatolian Neolithic component.

The slight but detectable Aegean signature in ROC001 is particularly notable. Indeed, given her young age, she was certainly part of the non-belligerent population of the site, and likely locally born, as also suggested by Sr isotope findings [9]. Our findings are in agreement with previous archaeological research showing intense trade relations between Roca Vecchia and the contemporary Middle Bronze Age Aegean world [5,15] and supposing that foreigners coming from the Minoic world, either Crete or mainland Greece, were living in Roca Vecchia and producing Aegean style pottery with local clay [2]. It is therefore plausible that a small nucleus of Aegean “foreigners” lived in Roca Vecchia in the form of a community colony [4] and interbred with locals, contributing to the particular genetic make-up of ROC001 (US2616 D).

## Methods

### Generation and Authentication of aDNA data

Archaeological samples were excavated by Prof. Cosimo Pagliara (formerly employed at the University of Salento), who held a permit for study, sampling and analysis of the samples from the concerned Soprintendenza. The samples were then passed on to PFF, director of the Anthropology lab after Prof. Pagliara’s retirement.

Sampling, surface decontamination, DNA extraction and library preparation were performed at the state-of-the-art ancient DNA facility of the Institute of Genomics, University of Tartu (Estonia), following protocols from Keller et. al, 2023 [24–26]. Four petrous bones and two teeth were sampled and then cleaned using dental tools to reduce surface contamination. We rinsed the samples using NaOCl 6%, Ethanol 70%, and MilliQ water (Millipore). After the rinsing, samples were left under UV lights for 1 hour and a half. Cell lysis and DNA extraction were performed by adding 0,5M EDTA (pH=8) (20x EDTA [μl] of sample weight [mg]) and Proteinase K (18,2 mg/mL) (0.5x Proteinase K [ml] of sample weight [mg]) for 72 hours at room temperature on a nutating mixer.

After three days, lysates were centrifuged to spin down pellet and bone chunks for 5 minutes at 4,000 rpm. The supernatant was collected and concentrated in Vivaspin 15-30kDa concentrators (Sartorius) for 25-45 minutes at 4,000 rpm. The purification steps were performed using the High Pure Viral Nucleic Acid Large Volume Kit columns (Roche). Ten times more Binding buffer (PB buffer (Qiagen)) than the sample volume was added and the concentrated samples were added to a 50 ml Roche column. The Roche columns were spun down at 4,000 rpm for 1 minute. DNA was washed using an Ethanol-based buffer (PE buffer (Qiagen)) to remove unbound substances. 100 µl EB buffer (Qiagen) was added to each sample and the columns were incubated for 10 minutes at 37°C to elute the DNA.

Starting from 30 µl of purified DNA solution, we built double indexed-double stranded DNA Illumina libraries using a non-UDG treatment. Library preparation included three stages: End Repair reaction, Adapter Ligase reaction and Adapter Fill-in reaction. Regarding the End Repair reaction, a 20 μl reaction mixture was prepared (per sample), containing 5 μl 10x End Repair Reaction Buffer (NEBNext), 2.5 μl End Repair Enzyme Mix (4 U/μl T4 DNA Polymerase, 10 U/μl T4 Polynucleotide Kinase) (NEBNext) and PCR grade water.

For the Adapter Ligase reaction, a 20 μl reaction mixture was prepared (per sample), containing 10 μl of 5x Quick Ligation Reaction Buffer (NEBNext), 5 μl of 2000 U/μl Quick T4 DNA Ligase (NEBNext), and 5 μl of 2.5 μM adapters mixture (oligos from Metabion). The Adapter Fill-in reaction was finally implemented using 20 μl reaction mixture per sample, containing 5 μl 10x Thermopol Buffer (NEBNext), 5 μl 8 U/μl Bst DNA Polymerase, Large Fragments (NEBNext), 0.8 μl 4x 25mM dNTP-mixture (Thermo) and 12.2 μl of PCR grade water.

After the first two steps, samples were purified using the Minelute PCR Purification Kit (Qiagen).

To prepare the PCR mixture, 20 μl of 10x HGS Reaction buffer (Eurogentec), 20 μl of 25 mM MgCl2 solution (Eurogentec), 10 μl of 20 mg/ml Bovine Serum Albumin (Thermo), 4 μl of 4×10mM dNTPs mixture (Thermo), 4 μl 5 U/μl HGS Taq Diamond Polymerase (Eurogentec) and 84 μl PCR grade water were taken per sample. The reaction mixture was divided into 142 μl portions in PCR tubes. 4 μl of two different 10 μM indexed primers (NEBNext) were added to each tube. Then, 50 μl of the mixture from the Adapter Fill-in reaction was added to each tube, mixed with a pipette, and 100 μl of each mixture was transferred to another pre-labeled PCR tube. Amplification of DNA fragments was performed using PCR under the following conditions: 5 minutes at 94°C, followed by 30 seconds at 94°C, 30 seconds at 60°C, 30 seconds at 68°C cycled for 15 times, and 7 minutes at 72°C, ending the PCR amplification at 4°C. Libraries were then purified using the MinElute PCR Purification Kit (Qiagen). Samples were finally quantified with Qubit™ dsDNA HS Assay Kit (Thermo) using the manufacturer protocol and analysed by Agilent Technologies 2200 TapeStation system and sequenced with Illumina NextSeq500/550 platform (paired-end sequencing 75bp -NSQTM 500 Mid-Output Reagent cartridge v2 150 cycles). One individual, ROC001, was resequenced with NSQTM 500 High-Output Reagent Cartridge v2 150 cycles).

### Raw-data processing and reads mapping

After the sequencing, adapter sequences, indexes and poly-G tails were cut from the reads through the software cutadapt 1.11 [27], which was used to discard sequences shorter than 30 bp.

Reads were merged by FLASH [28], and then mapped against GRChr3 (hs37d5) using Burrows-Wheeler Aligner, BWA 0.7.12 [29]. After the alignment, sequences were converted into BAM format and samtools 1.3 [29] was used to keep just the sequences that mapped against the human reference genome. Afterwards, Picard 2.12 (https://broadinstitute.github.io/picard/index.html) merged data from different flow cell lanes and removed duplicates. GATK 3.5 [30] was used for the indel realignment. Finally, reads with a quality lower than 10 (MQ10) were filtered out using samtools 1.9 [29].

### aDNA authentification

It is known that DNA degrades over time and, for the purpose of aDNA authentication, it can be distinguished from modern DNA according to specific characteristics: a) aDNA is highly fragmented and b) it shows a high-frequency C → T substitution at 5’ ends of sequences, due to cytosine deamination. MapDamage2.0 [17] was used to determine C to T transition frequencies. Samtools 1.9 determined the number of final reads, average length, average coverage and average endogenous content.

### Genetic sex estimation

Genetic sex was determined by estimating the fraction of reads mapping to chromosome Y out of all the reads mapping either to X or Y chromosome [31].

### Contamination estimation

We used two contamination estimation methods, implemented in ANGSD, as in [32] to assess the average male X contamination, while mitochondrial contamination was computed using ContamMix [33].

### mtDNA haplogroup determination

VCF files were generated using bcftools 1.14 [34] and uploaded on HaploGrep2 for the determination of mtDNA haplogroup [35]. Results were visually checked by aligning mapped reads to the reference sequence using Samtools 1.19 [36]. The haplogroup assignment was confirmed using PhyloTree [37].

### Calling of variants

Autosomal and sex chromosomal-related variants were called with ANGSD-0.934 commands --doHaploCall --doCounts --doMajorMinor calling all the positions in the Lazaridis et al 2016 aDNA dataset [38,39]. The output was converted to the PLINK format and prepared for the merge (see below) using ANGSD-0.917 and PLINK 1.9. The number of autosomal SNPs was calculated using PLINK 1.9 (Table 1).

### Kinship analysis

The two individuals were tested for kinship using READv2 [40] in PLINK format. Because running READv2 on only two individuals may yield unreliable results, particularly with low-coverage genomes, we applied a user-defined normalization value, computed on a genetically similar population, following the READv2 manual. For this purpose, genomes from Daunian individuals, a genetically similar reference population, were used [18]. Kinship was calculated on Daunians from Ordona, Salapia and San Giovanni Rotondo sites (excluding two Medieval individuals), and the median value was extracted from the “*meansP0_AncientDNA_normalized*” output file (column: “*nonnormalized_p0*”). This value was then used to estimate kinship for ROC001 and ROC006 with the flag *--norm_value*. Analyses were performed with and without normalization.

### Data processing and population genetic analyses

We merged the genetic data from ROC001 and ROC006 with the “1240K” dataset (version 54.1.p1) of Allen Ancient DNA Resource (AADR) [41] using EIGENSOFT 8.0.0 [42] and PLINK 1.9 [43]. We retrieved the West Eurasian and North African modern genetic data to be used as a scaffold in the PCA and ADMIXTURE analyses and we refined the list of ancient samples by selecting those from geographical regions relevant to this study (Supplementary Tables 3 and 4). For analyses involving only ancient samples, such as f4 statistics and qpAdm (see below), we utilized the autosomal “1240K” SNP set, whereas the autosomal “HO” SNP set was employed for analyses that also included modern populations (e.g., PCA and ADMIXTURE).

We performed the Principal Components Analysis (PCA) by mapping the genetic variation of the ancient samples onto the principal components derived from the “HO” AADR modern samples using the program *smartpca* implemented in EIGENSOFT 8.0.0 with the parameters “lsqproject” and “shrinkmode” (the full list of ancient and modern samples used in this PCA and ADMIXTURE analyses is in Supplementary Tables 3 and 4, respectively). Given the lower SNP coverage of ROC006, we performed additional PCA analyses to assess the robustness of its placement. First, we repeated the projection using a reduced set of modern individuals present in both the 1240K and HO datasets and the 1240K SNPs, to maximize the overlap with the ancient samples (the list of modern samples from the 1240K is in Supplementary Table 4). Second, we ran an independent PCA using only the HO SNP panel to control for any scaffold-related bias.

Using the “HO” AADR modern set, we employed the ADMIXTURE software [20] to perform unsupervised Admixture analyses with K values ranging from 2 to 8, projecting the ancient individuals onto the ancestral component allele frequencies inferred from the modern individuals.

We used the program qpDstat (option f4mode) implemented in the software ADMIXTOOLS 7.0.2 [19] to conduct f4 statistics analyses. We chose the sample South_Africa_2000BP.SG as outgroup in the f4 configurations.

We relied on the qpWave/qpAdm framework implemented within the suite ADMIXTOOLS 7.0.2 to explore the genetic composition of Roca Vecchia individuals, alongside other ancient samples of interest. As a first step, we tested two sets of four left sources combined with a list of reference sources (right) using the options “allsnps = YES” and “inbreed = NO”. The first set of left sources comprised Caucasus Hunter-Gatherer (CHG: KK1_noUDG.SG and SATP_noUDG.SG), Western Hunter-Gatherer (WHG: Luxembourg_Loschbour_published.DG), Anatolia Neolithic (19 samples labelled “Turkey_N”) and Yamnaya (10 individuals labelled “Russia_Samara_EBA_Yamnaya”), while in the second set, we replaced the Caucasus Hunter-Gatherers with the Iranian Neolithic source (5 samples flagged as Iran_GanjDareh_N). For both sets, we used the following right populations: Russia_Ust_Ishim_HG_published.DG (which was always kept as the first of the list), Russia_Kostenki14, Russia_AfontovaGora3, Eastern Hunter-Gatherers (I0061 = Russia_HG_Karelia, I0124 = Russia_HG_Samara, I0211 = Russia_HG_Karelia, and UzOO77 = Russia_EHG), Spain_ElMiron, Belgium_UP_GoyetQ116_1_published_all, Levant_N (Israel_PPNB), Russia_MA1_HG.SG, Israel_Natufian_published, Czech_Vestonice16, Caucasus Hunter-Gatherers and Iranian Neolithic.

Then, to better disentangle the contribution of Steppe-related ancestry in our samples, we implemented both a distal and a proximal qpAdm model. In the distal model, we replaced the Yamnaya source with Eastern Hunter-Gatherers (EHG), using also Anatolia_Neolithic, CHG, and WHG as left sources. The right (outgroup) populations included Russia_Ust_Ishim_HG_published.DG, Russia_Kostenki14, Russia_AfontovaGora3, Villabruna, Israel_Natufian_published and Iranian Neolithic. In the proximal model, we used a European Chalcolithic population pre-dating the Steppe expansion (Czech_C_Baalberge) and Yamnaya as left sources. The outgroup list was expanded to include all previously used ancient sources: Anatolia_Neolithic, CHG, EHG, WHG, Russia_Ust_Ishim_HG_published.DG, Russia_Kostenki14, Russia_AfontovaGora3, Villabruna, Israel_Natufian_published, and Iran_GanjDareh_N. For all qpAdm models, we considered results to be reliable only when the model yielded a p-value greater than 0.05 and all ancestry proportions were non-negative.

We inferred the ancestral composition of the two samples from Roca Vecchia, together with other Italian and West Eurasian individuals using PANE, which leverages non-negative least squares (NNLS) regression of the first 100 Principal components estimated as described above [44]. We used the following putative sources, averaging PC scores across individuals when possible: Caucasus Hunter-Gatherers (KK1_noUDG.SG), Western Hunter-Gatherer (Luxembourg_Loschbour_published.DG), Anatolia Neolithic (33 samples labelled “Turkey_N”) and Yamnaya (4 individuals labelled “Ukraine_EBA_Yamnaya”).

Graphical representations of the results were created using R 4.1.2 [45].

## Declarations

## Ethics approval and consent to participate

PFF holds permits from the relevant Soprintendenza to study and analyse the samples included in the current manuscript.

## Availability of data and materials

The DNA sequences generated during this study are available at the European Nucleotide Archive (ENA) at the accession number PRJEB93761. The data are also available in PLINK format through the data depository of the EBC-IG (https://evolbio.ut.ee/Aneli_2025/).

## Authors’contributions

L.P, P.F.F. designed the study; S.A., V.N., G.V, F.M., P.F.F and L.P wrote the manuscript; S.A., V.N, F.M and L.P ran population genomics analyses; V.N., S.S., T.S., H.K., A.S., and K.T. generated aDNA sequencing data; R.G, G.V. and P.F.F. ran anthropological and osteological analyses; All authors revised the manuscript.

## Acknowledgements

We thank C3S (http://c3s.unito.it) for providing the computational resources and Dr. Tuuli Reisberg and Dr. Reidar Andreson for supporting us with the sequences and genotypes upload process.

