## Supplementary material for "Ancient genomes from the siege and destruction of Middle Bronze Age Roca Vecchia (Apulia, Italy) shed light on Aegean contacts and conflicts": Supplementary Materials.pdf

**This PDF file includes:**

- Supplementary Note (Archaeological findings).
- Supplementary Figures related to the genetic analyses (from 7 to 11).

**Other Supplementary Materials for this manuscript include:**

- Supplementary Table 1. Archaeological and ancient DNA data of the individuals under study.
- Supplementary Table 2. READv2 results on kinship estimation.
- Supplementary Table 3. List of ancient samples used to run the PCA/PANE, Admixture, qpADM and f4 analyses.

- Supplementary Table 4. List of modern samples used as a scaffold for the PCA and Admixture analyses. Samples from the "HO" AADR dataset were used for the PCAs in Figure 2A and Supplementary Figure 8, while samples from the "1240K" were used for the PCAs in Supplementary Figures 9 and 10.
- Supplementary Table 5. f4 tests run on the newly sequenced ROC001 and ROC006 Roca Vecchia samples joined together.
- Supplementary Table 6. qpADM results for models including Iran Neolithic (Iran_GanjDareh_N), Western Hunter-Gatherer (Luxembourg_Loschbour_published.DG), Anatolia Neolithic (Turkey_N) and Yamnaya as sources (see Methods).
- Supplementary Table 7. qpADM results for models including Caucasus Hunter-Gatherer (CHG), Western Hunter-Gatherer (Luxembourg_Loschbour_published.DG), Anatolia Neolithic (Turkey_N) and Yamnaya as sources (see Methods).
- Supplementary Table 8. qpADM results for a "distant" model including Anatolia Neolithic (Turkey_N), Western Hunter-Gatherer (Luxembourg_Loschbour_published.DG), Caucasus Hunter-Gatherer (CHG) and Eastern Hunter-Gatherer as sources (see Methods).
- Supplementary Table 9. qpADM results for a "proximal" model including European Chalcolithic (Czech_C_Baalberge), and Yamnaya as sources (see Methods).
- Supplementary Table 10. Results of the PANE analyses reporting the contribution of Caucasus Hunter-Gatherer (Georgia_Kotias.SG), Western Hunter-Gatherer (Luxembourg_Loschbour), Anatolia Neolithic (Turkey_N) and Yamnaya (Ukraine_EBA_Yamnaya) to Italian (sheet "Italy") and Eurasian (sheet "Eurasia") populations.

### Supplementary Note

#### Archaeological setting

The powerful Middle Bronze Age wall at Roca Vecchia (Lecce, Italy) is preserved for a length of about 200 m, it was surrounded, along its external front, by a ditch dug in the calcarenite. A centrally placed monumental main gate and five long and narrow posterns (named A through E) have been found so far (Guglielmino, 2005; Scarano, 2012; Guglielmino and Pagliara, 2017) (Supplementary Figure 1). The fortification and the site were destroyed, between the half of the XV century and the beginning of the XIV BC by a large fire-event. At the moment of destruction the outer access of at least three posterns, B, C and D, were closed by wooden or stone walls (Supplementary Figure 2). In the obstructed posterns hundreds of vases of various size and form, stoves, bone and metal tools, clay spindle whorls, animal bones, traces of hearths and others items have been unearthed. These finds are typical of dwelling contexts and they suggest that, once obstructed, the posterns were used as shelters by part of the population. It seems likely that the posterns were used only during peaceful times and were blocked during war times as they would weaken the fortification. The custom of obstructing the posterns during war times has been supposed in the nearly coeval site of Coppa Nevigata (Recchia 2008) and in many archaeological sites across the Mediterranean. As far as Bronze Age sites are concerned, it is possible to recall et-Tell, a site supposed to be the biblical town of Ai, where a postern of the Ancient Bronze II fortification was blocked by a wall, probably during a siege (Callaway 1976). The reuse of posterns as dwelling areas is probably due to the overpopulation of the sites during war times when people living in the surroundings found shelter inside the fortification, a similar behaviour has been hypothesized for the Mycenaean citadels and for Troy (Iakovidis 1983; Mylonas 1964). It is most likely that the obstruction of the posterns and the huge fire that eventually destroyed Roca Vecchia are the result of a siege that took place during Middle Bronze Age 3 (MBA3), specifically between the second half of the 15th century BCE and the first decades of the 14$^{th}$ century BCE (Guglielmino and Pagliara, 2007, Calcagnile et al., 2012, Scarano, 2012). Nine human skeletons have been found under the debris of the fortifications. The postern C has a preserved length of 15 m, a maximum breadth and height of 1.5 m, it is oriented NE (inner entrance)-SW (outer obstructed entrance). At the bottom of the layer infilling the postern, labelled US 2616, close to its obstructed outer entrance and resting on the floor, seven human skeletons (Fabbri, 2020), labelled US 2616 A to G, comprising both adult and immature individuals of various ages, were found. The Main Gate is preserved for a length of 22 m, with a maximum breadth of 3.5 m and maximum height of 4 m, it was filled with the debris of the collapsed fortifications comprising huge amount of ash and charred logs. Two human adult skeletons have been found: one (Vincenti et al., 2024), labelled US 813 RA2, in a side room, labelled Room B; another (Maggio et al., 2020), towards the inner entrance, labelled US 14203 RA7.

### Anthropological assessment of the remains

#### Postern B

The child skeleton found at the base of the infilling of postern B is still unpublished, his/her dental age (Alqahtani et al., 2011) is estimated to 1y±4m.

#### Postern C

Data concerning the individuals from postern C are resumed from a paper, Fabbri, 2002, and from an unpublished communication, Fabbri, 2020. At the base of US 2616 seven human skeletons (labelled US 2616 A through US 2616 G) have been found (Supplementary Figure 1). The skeletons were in a limited area of about 2x1.5m edged by the wall obstructing the outer entrance and by a row of big vases (height 80cm) which run perpendicularly to the main axe of the postern. Two of the individuals (Supplementary Figure 2), US 2616 A and US 2616 B, were adults, one, US 2616 C, was an adolescent and four, US 2616 D, US 2616 E, US 2616 F and US 2616 G, were children. According to pelvis-based sex determination (Phenice 1969), US 2616 A was a male. For US 2616 B, whose pelvic bones are very fragmented, we considered cranial features (Ferembach et al., 1979) indicating that female sex is probable for this individual, the comparison of the two adult skeletons shows that US 2616 B is evidently shorter and more gracile than US 2616 A, thus confirming the proposed female sex determination. US 2616 A's partially preserved right pubic symphysis shows no billowing and a clear dorsal rim. Following Meindl et al. (1985) the age should be over 30 (absence of billowing) and probably over 40 (presence of dorsal rim). US 2616 B's right hip bone preserves the majority of its auricular surface, features proposed by Buckberry and Chamberlain (2002) can be estimated and correspond to the "auricular surface stage II" observed at a mean age of 29,33 years (median 27 years, range 21-38 years). In US 2616 C the preserved long bone extremities are not fused to the bone shafts, indicating an age lower than 17-18 years. The partial fusion of the secondary ossification centers of the bones of hands and feet, which takes place between 16 and 20 years of age (Ferembach et al., 1979), suggests that the most probable age for this individual is 16-17 years. As regards the four children, their ages have been determined considering the degree of maturation and eruption of the dentition (Alqahtani et al., 2011) and the lengths of long bones (Stloukal and Hanáková, 1978). Their most probable ages at death are: 8-10 years for US 2616 D; around 4 years for US 2616 E; 7-8 years for US 2616 F; around 7 years for US 2616 G. No traces of *perimortem* traumas have been observed on the seven skeletons from postern C.

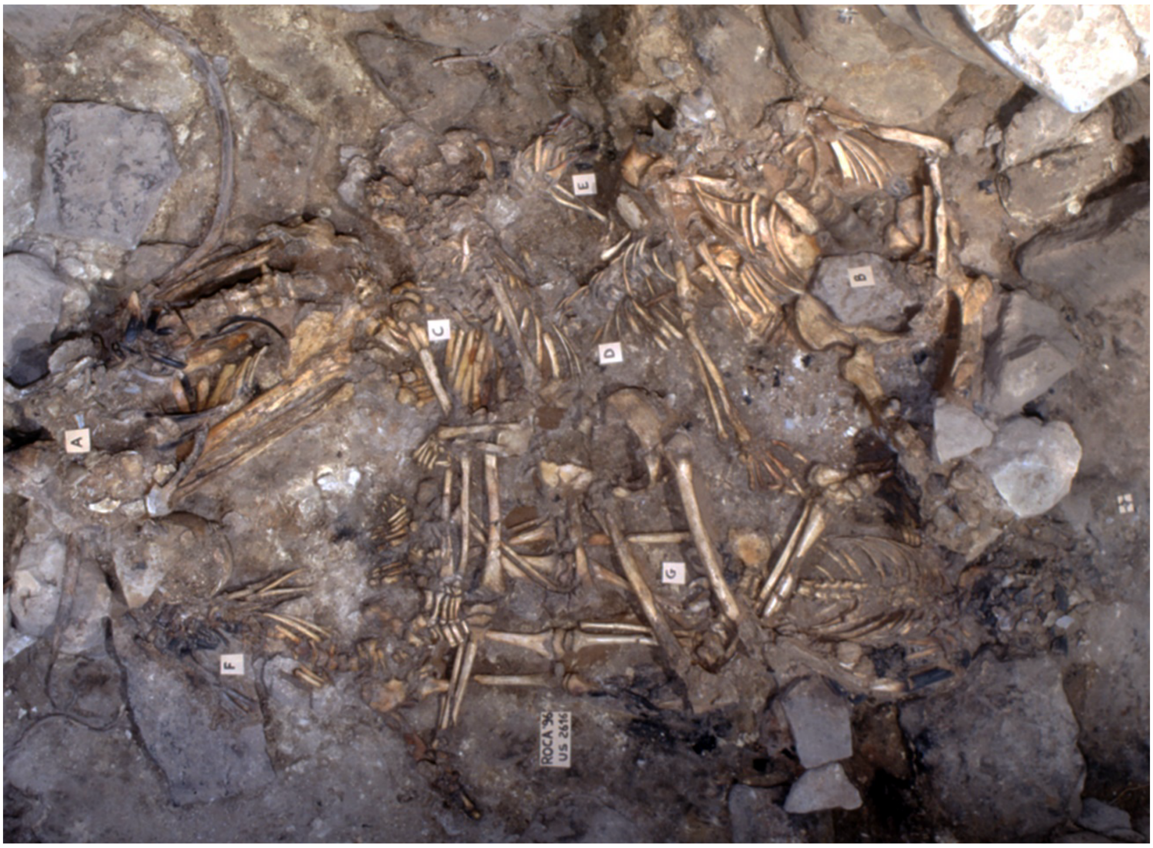


**Supplementary Figure 1.** The seven human skeletons found in postern C (US 2616 A-G).

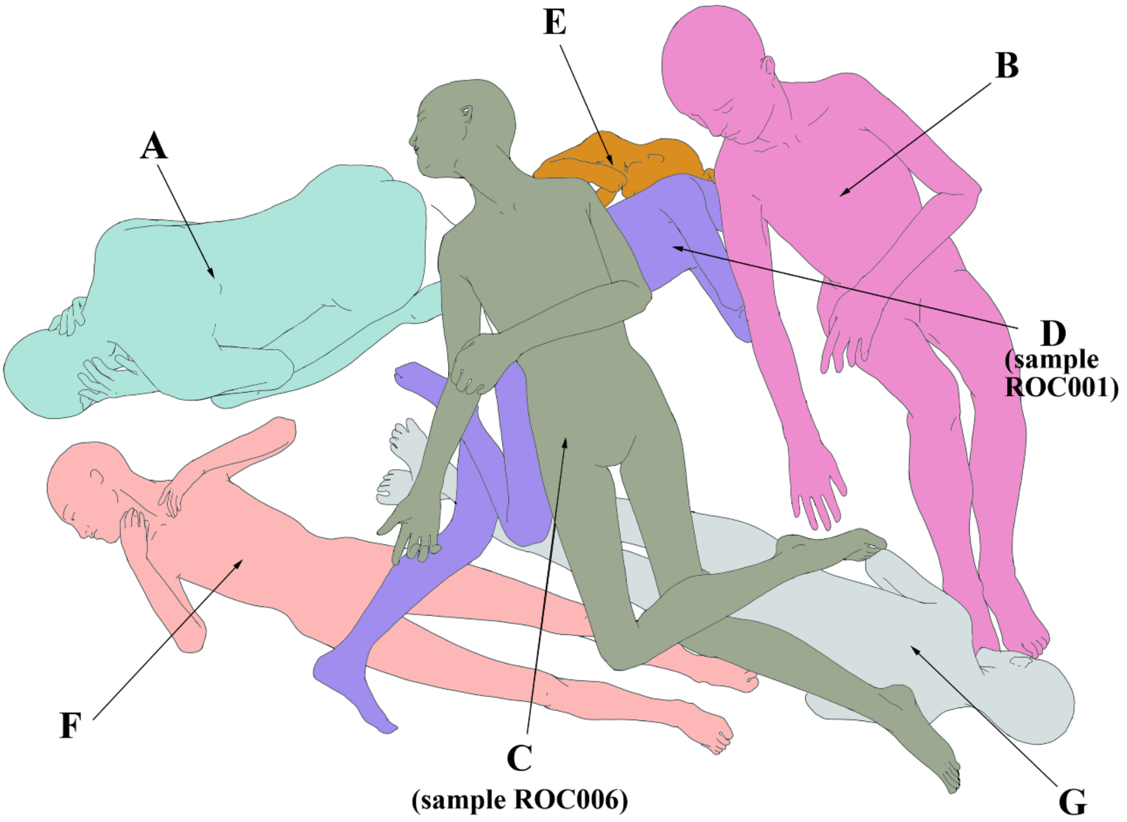


**Supplementary Figure 2.** Reconstruction of the positions of the bodies of the seven human skeletons (US 2616 A-G) found in postern C, with indication of the two individuals whose aDNA was retrieved.

**The Main gate**

The human skeleton US 14203 RA7 (Supplementary Figures 3 and 4) was in very bad state of preservation, following Maggio et al., 2020 cranial features, Acsadi and Nemeskeri formula, and postcranial measures, particularly humeral diaphyseal ones are on the male side of the variation; as to age at death, dental wear (grade 4-5 on preserved teeth, Molnar, 1971), ante-mortem loss of all mandibular molars and marked retraction of the mandibular alveolar border suggest that US 1402 RA7 was an adult of medium or advanced age, most probably more than 40-year-old. Maggio et al., 2020 observed the presence of a depressed blunt force trauma on the left hemifrontal, traces of osteoblastic and osteoclastic activities on the trauma show that it is *antemortem* and that it took place at least two months before his death.

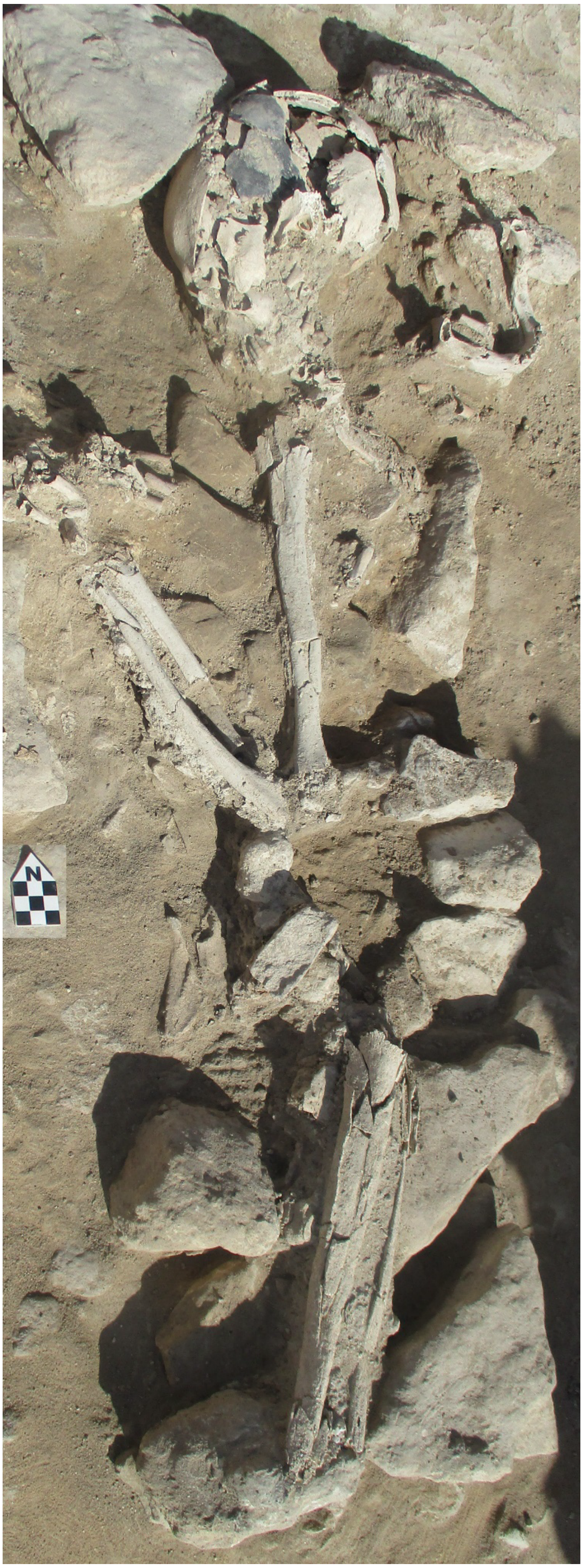


**Supplementary Figure 3.** The human skeleton found at the inner entrance of the Main Gate (US 14203 RA7).

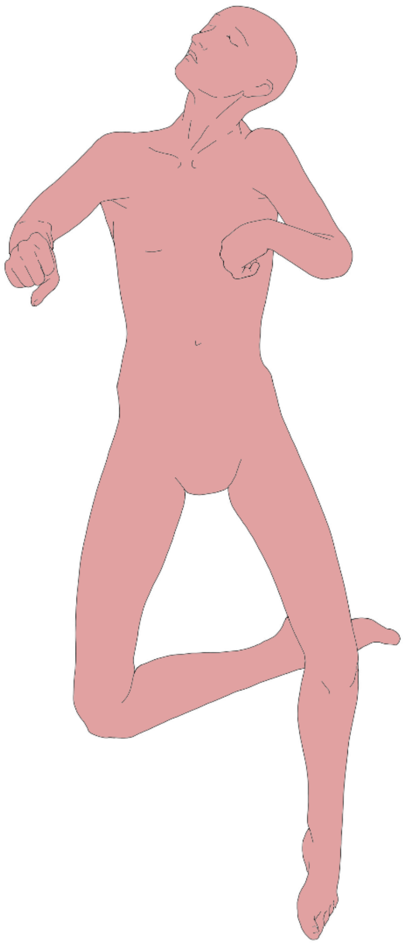

**Supplementary Figure 4.** Reconstruction of the positions of the body of the human skeleton found at the inner entrance of the Main Gate (US 14203 RA7).

Following Vincenti et al., 2024, (Supplementary Figures 5 and 6) the human skeleton found in room B of the main gate, US813 RA2, was a young male according to pelvis features (Phenice, 1969; Walker, 2005), he was aged approximately 18–19 years (Cameriere et al., 2009). At least four *perimortem* sharp force traumas were present on the bones of US 813 RA2: one on the 3rd right rib and T4; two on the 11th and 12th right ribs; and one on the 12th left rib and L2. All injuries were due to blows inflicted from behind and seem to be the result of close combat. The blows could have been inflicted by two different aggressors, one of whom probably left-handed, or by a single person brandishing two weapons simultaneously. This particular type of combat is attested in contemporary Aegean iconography (Vincenti et al., 2024).

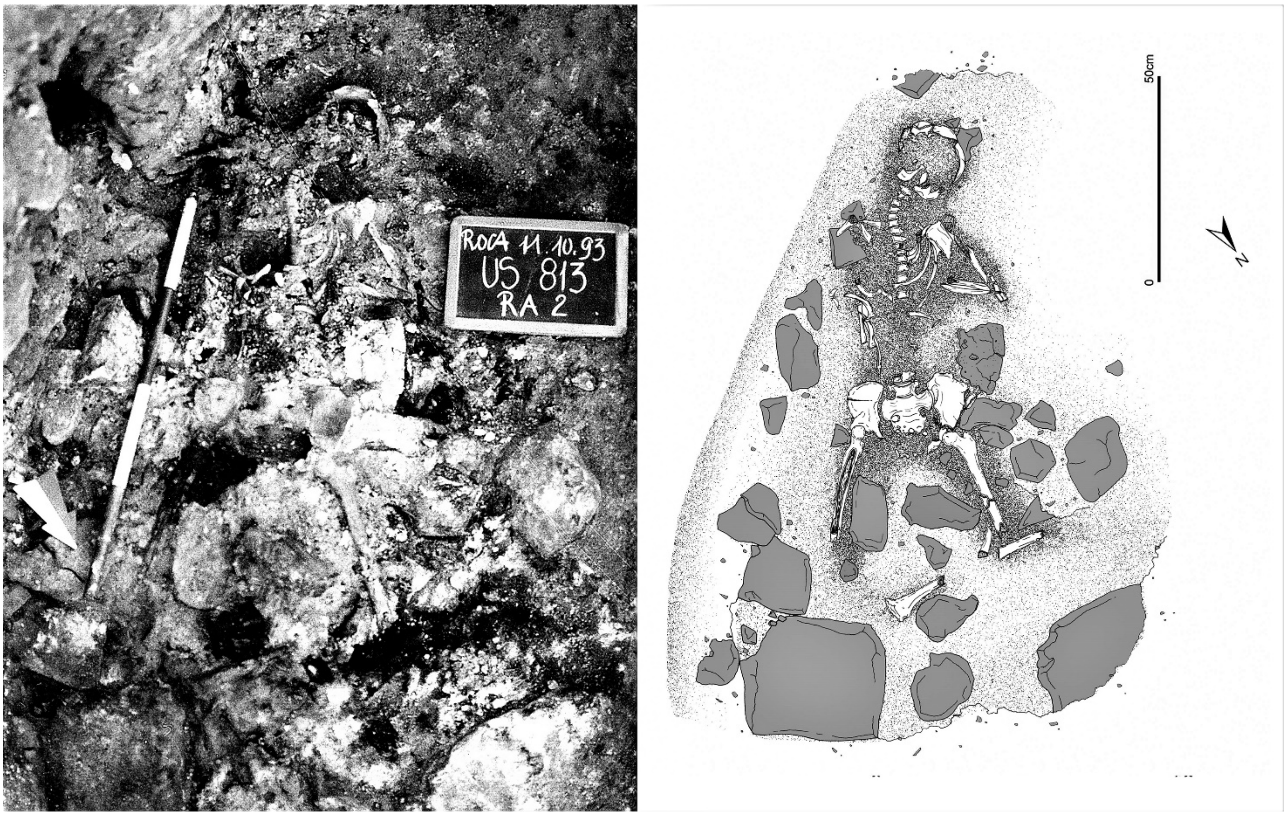


**Supplementary Figure 5.** The human skeleton found in the room B of the Main Gate (US 813 R2).

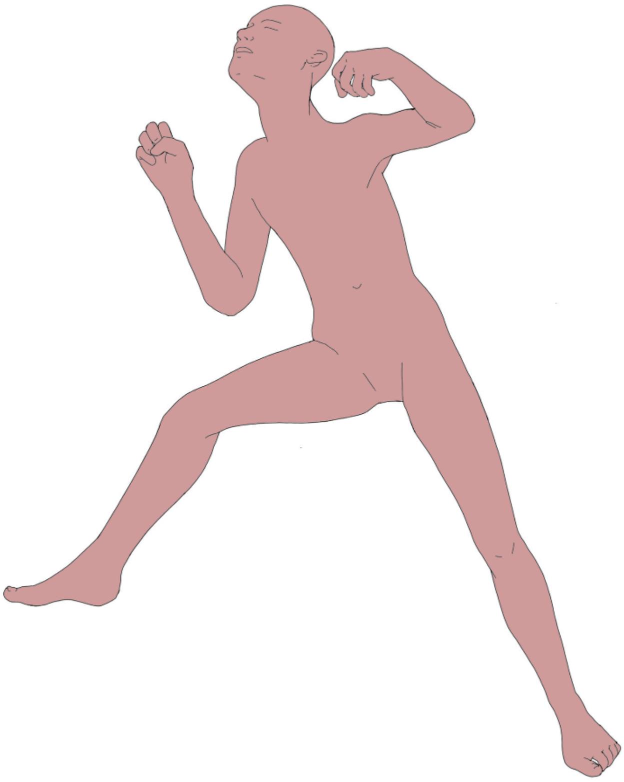

**Supplementary Figure 6.** Reconstruction of the positions of the body of the human skeleton found in the room B of the Main Gate (US 813 R2).

All the aforementioned individuals were placed beneath a huge stratum, reaching 4 m in the area of the main gate, of debris containing burned stones, charred logs and abundant ashes and charcoals, which was formed by the collapse of the overhanging structures of the fortification. The coherent stratum of tens of cubic meters allows to exclude that the human remains were intentionally buried, moreover skeleton postures and the lack of associated burial goods would be unmatched among local contemporary burials (Orlando, 1995).

Many of the bones of the nine published individuals showed traces of *perimortem* heat exposure and some of them (US 14203 RA7, US 813 RA2 US 2616B, US 2616C and perhaps US 2616 F) have been found in a position compatible with the pugilistic posture (Fabbri, 2002, 2020, Maggio et al., 2020, Vincenti et al., 2024). The pugilistic posture is characterized by head hyperextension, hands closed into fists, flexed toes, extreme limb flexion, arms abducted at the shoulder joint and flexion at the elbow joint, lower limb abduction at the hip joint, knee flexion, and foot extension (Bohnert, 2004, Symes et al., 2008). The pugilistic posture is assumed by human bodies exposed to heat for at least 10–20 min. The bones were fragmented, with both heat-induced fractures and crush fractures due to the collapse and weight of overhanging structures. Particularly important for establishing the pre-combustion state of the remains are heat-induced fractures and warping. Bone heat-induced fractures caused by collagen removal, which reduces elasticity by changing resistance to traction, the structural integrity of the bone tissue is compromised and fractures along the lines of greater stress are formed (Ortner and Putschar, 1981; Herrmann and Bennett, 1999): in particular, the curved transverse fractures (perpendicular to the diaphysis axis of long bones and appear arched) are found primarily, but not exclusively, in freshly burned bones (fleshed or defleshed), warping occurs only in a small percentage of dry bones (about 8% of the examined remains; Gonçalves et al., 2015). Individual US 2616 A, B and C present transversal curve fractures, US 2616 F and E warping. Individual US 813 RA2 bears curved transverse fractures and warping. Individual US 14203 RA7 showed only limited signs of heat exposure: no heat-induced fractures were recognizable, and color changes related to thermal exposure were present in a limited area of the skull.

The huge fire which destroyed the site of Roca Vecchia during Middle Bronze Age clearly appears as the final outcome of a prolonged siege. During the siege, the external ends of at least three posterns were obstructed and they were transformed in dwelling places, a common behaviour in fortified sites during wartimes. The finding of at least ten unburied human skeletons reinforces the interpretation that warfare took place in Roca Vecchia (Walker, 2001, Martin and Harrod, 2015, Erdal, 2012). Among the human individuals found under the debris of the fortifications, one, US 813 RA2 found in Room B of the main gate, was a belligerent who was stabbed to death and could have been as well an attacker as a defender, the seven from Postern C, US 2616 A to G, were non belligerent and were part of the defending population, the last one, US 14203 RA7 from the main gate, was probably an attacker or defender belligerent.

### Other Supplementary figures

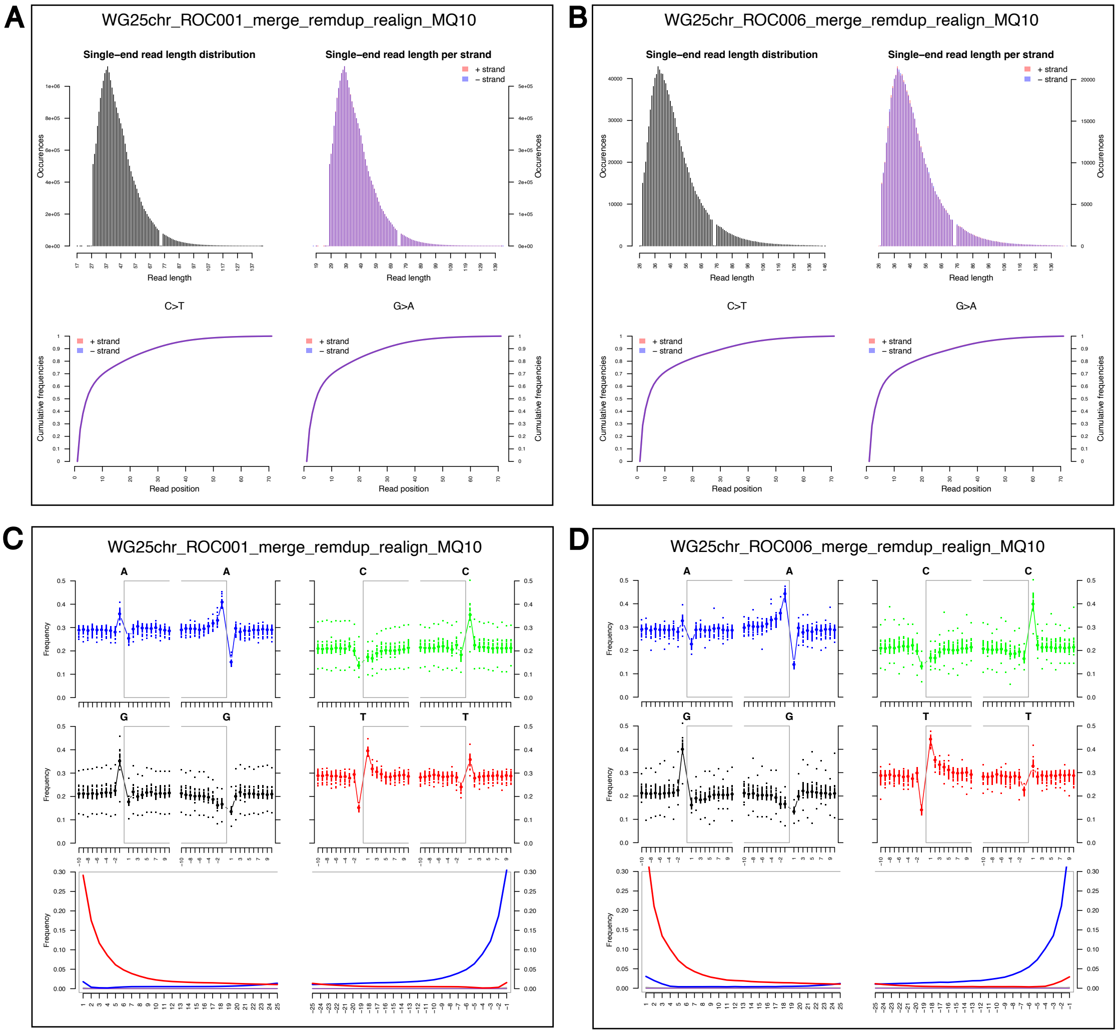


**Supplementary Figure 7.** Fragment length, misincorporation and damage analyses performed by MapDamage2. In panels A and B, the upper histograms on the left show the overall distribution of sequenced fragments, while the histograms on the right display the same distribution, distinguishing reads mapped to positive and negative strands of DNA. The lower panels show the cumulative frequencies for C to T and G to A substitutions, according to the position along the read length, scaled based on the first 70 positions. In panels C and D, the upper panel shows the base-specific frequencies within the read length (delimited by the light grey square brackets) and outside the read length. Typical aDNA damage signatures for authenticity are quite evident in the green and black plots, showing C to T (5' end) and G to A (3' end) substitutions, respectively. The lower panels show substitution frequencies at specific position from 5' and 3' ends, with red indicating C to T substitution distributions and blue indicating G to A substitution distribution, along the read length. These patterns represent another, typical molecular signature of ancient DNA.

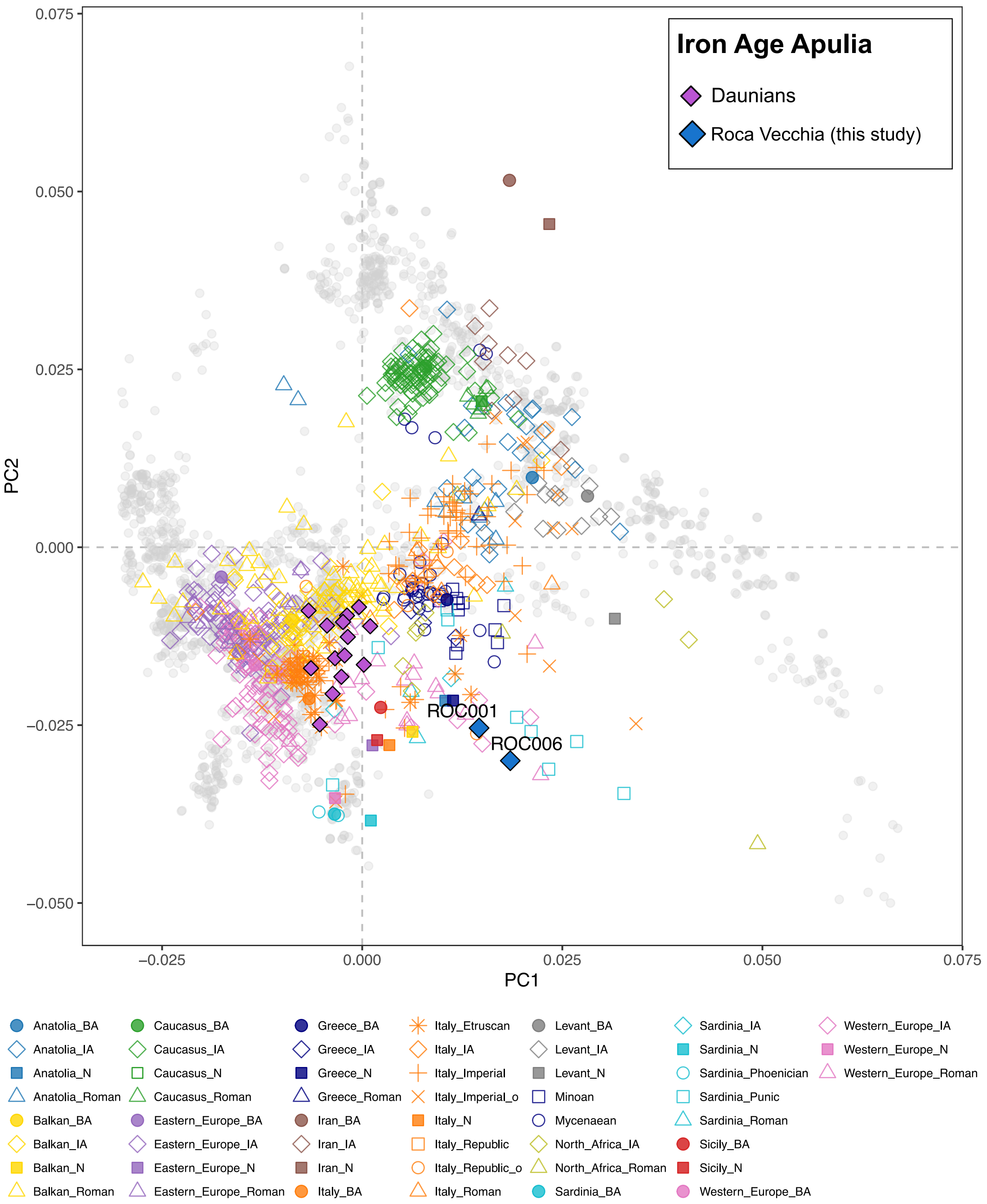


**Supplementary Figure 8.** Principal Component Analysis (PCA) showing the two newly reported individuals (ROC001 and ROC006) in the context of available ancient DNA from the region (Supplementary Table 3). For Neolithic and Bronze Age groups, only average positions are plotted. A more focused version of this PCA, showing only Italian Iron Age samples as reference, is provided in Figure 2A. The PCA was performed using the "HO" dataset of the Allen Ancient DNA Resource as a modern reference scaffold (AADR; see the Methods section and Supplementary Table 4).

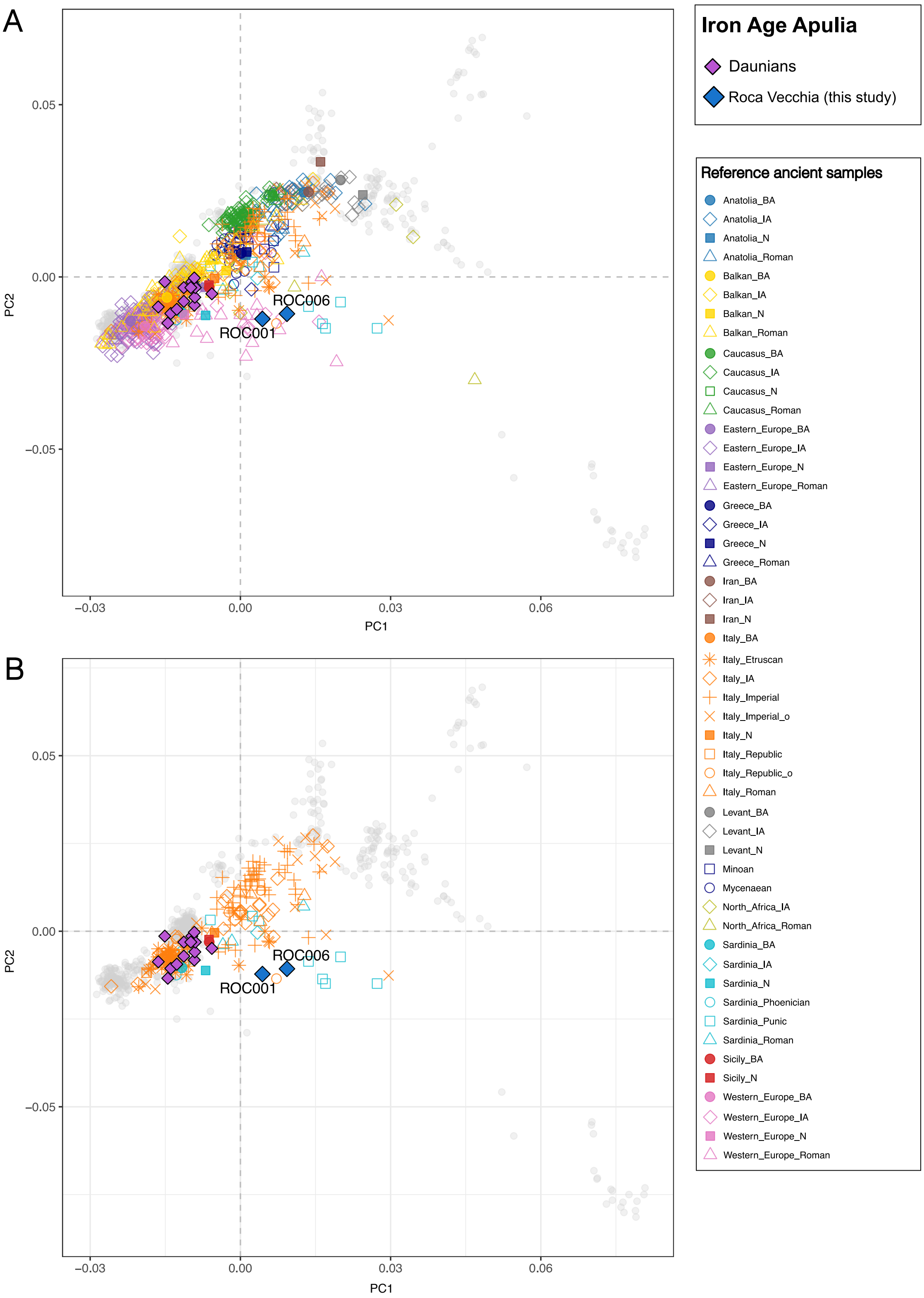


**Supplementary Figure 9.** Principal Component Analysis (PCA) showing the genetic projection of the two newly reported individuals (ROC001 and ROC006) onto a modern reference scaffold constructed from the "1240K" dataset of the Allen Ancient DNA Resource (AADR; see the Methods section and Supplementary Table 4). Panel A includes a selection of the ancient individuals from across Europe and the Mediterranean region, while Panel B displays only those from the Italian Peninsula.

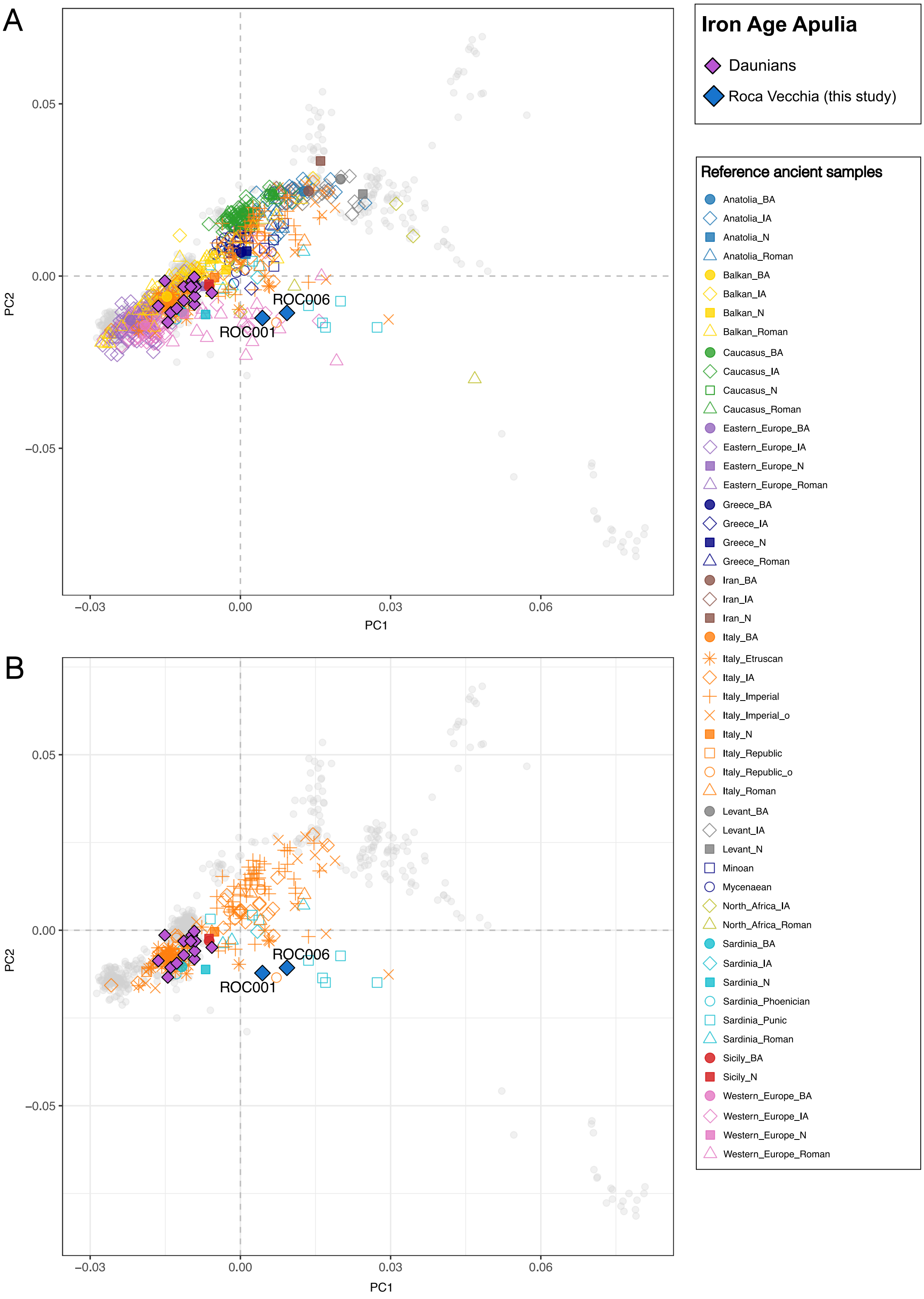


**Supplementary Figure 10.** Principal Component Analysis (PCA) showing the genetic projection of the two newly reported individuals (ROC001 and ROC006) onto a modern reference scaffold constructed from the "1240K" dataset of the Allen Ancient DNA Resource (AADR; see the Methods section and Supplementary Table 4), using SNPs from the "HO" panel. This analysis was performed to reinforce the placement of individual ROC006, which has a lower number of called genetic positions, within the PCA space (Figure 2A and Supplementary Figures 8).

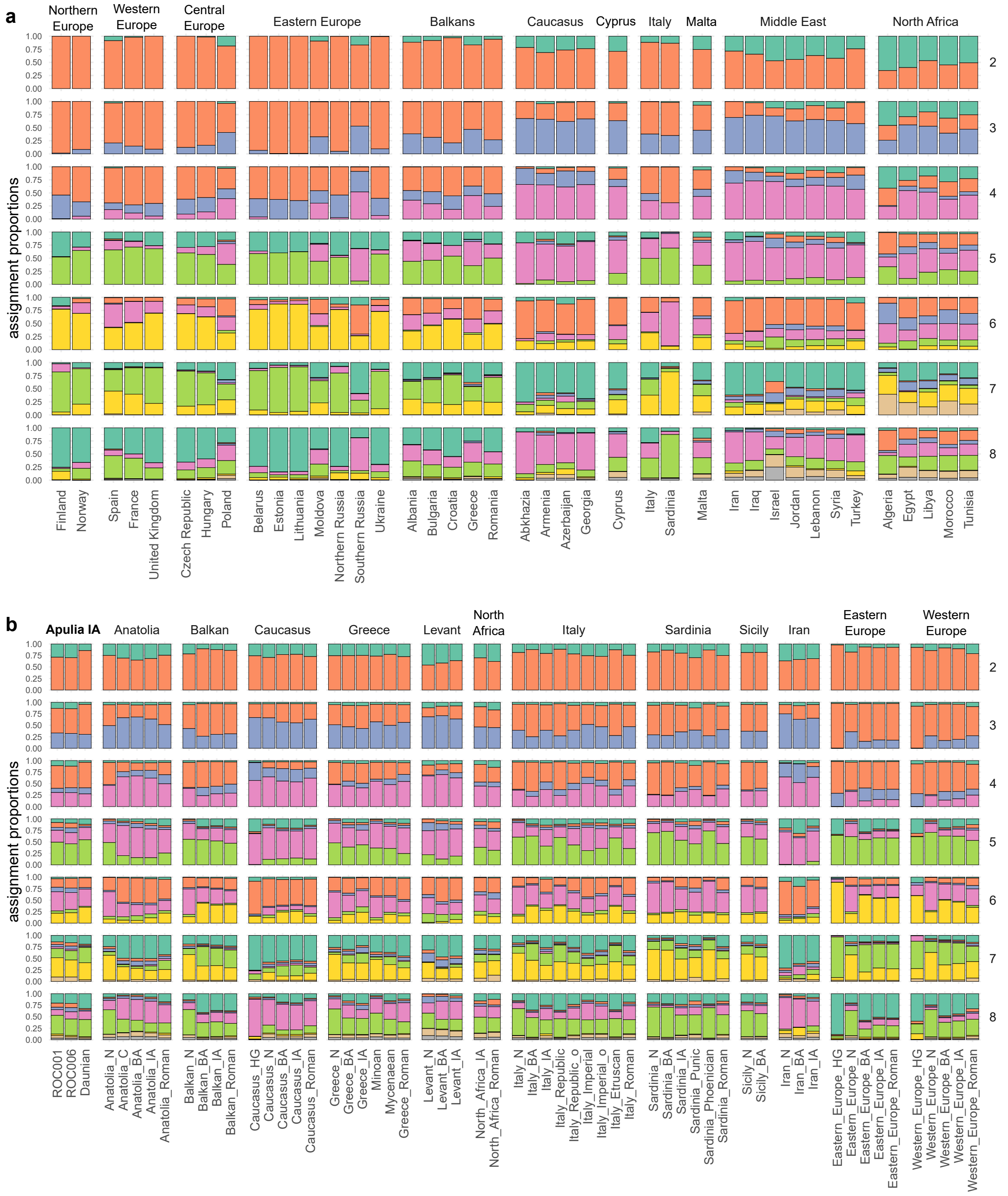


**Supplementary Figure 11.** Full Admixture plot showing samples used to infer the admixture space (a) and samples projected onto the inferred space (b).
